# Application of ^2^H_2_O to quantify red blood cell protein synthesis rates in young trained and untrained males and females

**DOI:** 10.1101/2024.10.28.620651

**Authors:** Hilkka Kontro, Chris McGlory, Martin J MacInnis

## Abstract

**Aim:** Compare the fractional synthetic rate (FSR) of hemoglobin in trained and untrained humans (Hb FSR).

**Methods:** We employed deuterated water (^2^H_2_O) to measure Hb FSR in young males (n=10) and females (n=10) who were aerobically trained (n=5 per sex) and untrained (n=5 per sex). Overall, participants had a mean 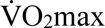 of 49.8 [SD: 10.9] mL/kg/min, hemoglobin mass of 775 [180] g, and red blood cell volume of 2370 [550] mL. After an initial loading dose, participants ingested ^2^H_2_O daily for 28 days to maintain a stable ^2^H body water enrichment (∼0.9 atom percent deuterium (APD)), as measured in saliva samples collected every 2-3 days. ^2^H-enriched alanine was measured in RBC protein using gas chromatography combustion isotope ratio mass spectrometry.

**Results:** The increase in APD for Hb protein was nonlinear for the first two weeks, but stabilized from day 14 to day 28, with mean APD reaching 0.033 [0.005] % on day 28. Hb FSR calculated over this 2-week period was 0.84 [0.15] %/day, which equated to an Hb absolute synthetic rate of 6.5 [2.2] g/day and a lifespan of 126 [30] days. Hb FSR was not different trained (0.83 [0.19] %/day) and untrained (0.86 [0.24] %/day, P=0.81) individuals or between males (0.80 [0.25] %/day) and females (0.88 [0.17] %/day), P=0.24).

**Conclusion:** Habitual endurance training does not appear to affect Hb FSR, but the use of ^2^H_2_O to measure Hb FSR in humans has potential applications to many experimental and clinical scenarios.

## INTRODUCTION

Comprising ∼97% of all red blood cell (RBC) proteins, hemoglobin (Hb) is a heme protein that binds and transports oxygen throughout the body (Weed et al., 1963). There are no direct methods to assess the rate at which Hb (or RBCs) are produced. Existing methods, which are typically complex, measure the rate of RBC destruction and infer a rate of synthesis; however, characteristics of Hb make it feasible to assess the fractional synthetic rate (FSR) of RBC proteins with the deuterated water (^2^H_2_O) method (Gasier et al., 2010). As almost all of the proteins in RBCs are synthesized during maturation in the bone marrow (Palis, 2014), there is a delay between the synthesis of Hb and its appearance in the blood. Nevertheless, Hb FSR can theoretically be measured when a linear increase in alanine-bound ^2^H isolated from Hb is observed in conjunction with stable ^2^H body water enrichment, albeit an extended protocol (i.e., several weeks) is likely required to account for the delay between protein synthesis and appearance in circulation. Although the precursor pool (i.e., bone marrow) is difficult to access in humans, measuring body water ^2^H enrichment is non-invasive and straightforward.

Furthermore, because the total mass of Hb in circulation (i.e., Hb_mass_) can be measured in humans (Burge & Skinner, 1995), it is also theoretically possible to measure the Hb absolute synthetic rate (ASR). A ^2^H_2_O technique for assessing Hb FSR would facilitate investigations into various aspects of human physiology.

Measuring Hb FSR could provide insights into exercise training adaptations. Chronic endurance training is associated with an elevated blood volume and hemoglobin mass (Hb_mass_) (Heinicke et al., 2001; Kjellberg et al., 1949; Montero et al., 2017; Schmidt & Prommer, 2008) that contributes to enhanced maximal aerobic capacity 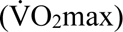 (Hagberg et al., 1998; Kanstrup & Ekblom, 1982). In addition, the RBC lifespan for endurance athletes is reported to be shorter than untrained individuals (Mairbäurl et al., 1983; Weight et al., 1991). The maintenance of a higher RBC volume with a shorter average cell life would necessitate a faster Hb turnover rate. A handful of studies have indicated that endurance athletes show signs of hyperplasia of the hematopoietic bone marrow (Caldemeyer et al., 1996; Shellock et al., 1992; Vogt et al., 2008) that may facilitate a higher Hb synthesis rate than in untrained humans.

The aims of the present study were to establish an oral stable isotope tracer method (i.e., ^2^H_2_O) to measure Hb FSR and ASR and measure potential differences in Hb FSR and ASR between habitually trained and untrained individuals. We hypothesized that Hb FSR and ASR could be measured using a 4-week protocol and that the Hb FSR and ASR would be higher in trained vs. untrained volunteers. We also sought to explore potential sex-related differences in blood protein synthesis rates, as well-established differences in Hb_mass_ and in iron status exist between the sexes (Finch & Cook, 1984; Goodrich et al., 2020; Kontro et al., 2024).

## METHODS

### Ethical Approval

All participants provided written, informed consent prior to enrollment in the study and were required to pass the Physical Activity Readiness Questionnaire (PAR-Q+). The study was approved by the University of Calgary Conjoint Health Research Ethics Board (REB19-0215) and conformed to the standards set by the Declaration of Helsinki, except for registration in a database.

### Study design

This cross-sectional study investigated Hb synthesis in healthy endurance-trained and untrained males and females during 4 weeks of habitual living (Figure 1). Participants completed an aerobic fitness test and a Hb_mass_ assessment prior to baseline sample collection. Thereafter, participants consumed ^2^H_2_O daily for 4 weeks. Regular blood and saliva samples were collected during this period, and Hb_mass_ was assessed again at the end of the study. Participants were required to maintain their typical training habits throughout the study and keep a training log of all their training activities. Albumin FSR was also assessed as an internal control.

**Figure 1.**
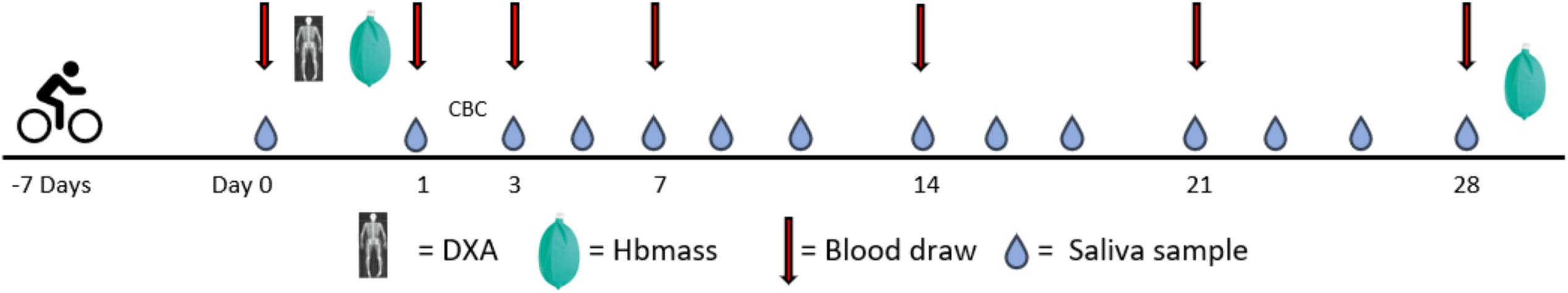
Timeline of measurements and sampling of blood and saliva. One dose of ^2^H_2_O was ingested daily after the loading phase of eight doses on Day 0.

The RBC lifespan was used as a proxy for Hb turnover for statistical power. Based on the results of Weight et al. (1991), where trained runners and untrained controls had RBC lifespans of ∼70 [25] days and 114 [30] days, we calculated an effect size of 1.6. The required sample size (α = 0.05 and 1 – β = 0.80) for this effect size was 8 participants per group (16 in total), and 20 were recruited to protect statistical power and account for participant attrition.

### Participants

Twenty volunteers (10 F, 10 M) partook in the study, with equal numbers of trained and untrained participants within each sex. Females and males were approximately matched for aerobic fitness when 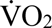 max was normalized to fat-free mass (Table 1; P=0.57). Over the preceding 3 months, self-reported habitual endurance training volume was 8.4 [2.9] h/week for the trained group and 1.2 [1.0] h/week for the untrained group. The trained participants had been involved in endurance sports for 10 [6] years. Participants’ primary sports were cycling (n=4), running (n=2), triathlon (n=2), cross-country skiing (n=1), and rowing (n=1). For the untrained group, an activity level corresponding to the minimum suggestion of the ASCM physical activity guidelines (∼150 min/week of moderate-vigorous exercise) or less was accepted with no recent history of competitive endurance sports. Both eumenorrheic females (n=4) and females on hormonal contraceptives (n=6) were included. No participants were amenorrheic.

**Table 1.** Participant characteristics at baseline disaggregated by sex and training status.

|  | All<br>(n=20) | Trained<br>(n=10) | Untrained<br>(n=10) | Males<br>(n=10) | Females<br>(n=10) | P value (Training Status x sex;<br>Training status; Sex) |
| --- | --- | --- | --- | --- | --- | --- |
| Age (y) | 28 [6] | 30 [6] | 26 [7] | 29 [8] | 27 [4] | 0.78; 0.28;0.34 |
| BM (kg) | 66.2 [10.9] | 65.8 [11.4] | 66.7 [11.0] | 73.7 [10.2] | 58.8 [3.1] | 0.39; 0.80; **0.001** |
| FFM (kg) | 51.3 [10.8] | 52.5 [11.3] | 50.1 [10.8] | 59.1 [10.1] | 43.5 [3.1] | 0.92; 0.50;<**0.001** |
| Hb<sub>mass</sub> (g) | 775 [180] | 815 [193] | 753 [166] | 902 [157] | 648 [89] | 0.96; 0.18;<**0.001** |
| Hb<sub>mass</sub> (g·kg<sup>-1</sup>) | 11.6 [1.4] | 12.3 [1.4] | 10.9 [1.0] | 12.2 [1.5] | 11.0 [1.0] | 0.33; **0.008**; **0.013** |
| Hb<sub>mass</sub> (g·kg FFM<sup>-1</sup>) | 15.1 [1.7] | 15.6 [2.1] | 14.7 [1.2] | 15.4 [1.9] | 14.9 [1.5] | 0.94; 0.27; 0.54 |
| [Hb] (g·dL<sup>-1</sup>) | 15.2 [1.3] | 15.2 [1.3] | 15.5 [1.5] | 16.2 [1.0] | 14.2 [0.6] | 0.45; 0.09; <**0.001** |
| $\dot{V}O_2\text{max}$ (mL·kg<sup>-1</sup>·min<sup>-1</sup>) | 49.8 [10.9] | 58.8 [6.0] | 40.7 [5.4] | 53.1 [11.2] | 46.4 [10.0] | 0.47; <**0.001**; **0.006** |
| $\dot{V}O_2\text{max}$ (mL·kg FFM<sup>-1</sup>·min<sup>-1</sup>) | 64.3 [12.5] | 74.1 [8.3] | 54.4 [6.9] | 65.9 [11.5] | 62.6 [13.9] | 0.68; <**0.001**; 0.36 |
| Training volume (h·wk<sup>-1</sup>) | 4.9 [4.4] | 8.6 [3.0] | 1.2 [1.0] | 4.8 [4.4] | 5.0 [4.2] | 0.34; <**0.001**; 0.67 |

*Note:* BM = Body mass; FFM = fat-free mass; Hb = hemoglobin; Hb<sub>mass</sub> = Hemoglobin mass; $\dot{V}O_2\text{max}$ = maximal oxygen uptake; M
= male; F = female. Data are reported as mean (SD). Bold: P<0.05

### Data Collection

#### Training load

Participants wore a HR monitor (4iii, Cochrane, AB, Canada) and provided their session Rating of Perceived Exertion values (sRPE) for all exercise training sessions. The 4-week training load of the participants was assessed using total training volume and sRPE values (Foster et al., 2001).

#### Body composition

Whole-body dual-energy X-ray absorptiometry (DXA) scans by Lunar iDXA (GE Healthcare, Chicago, IL, USA) were used to quantify fat free mass (FFM).

#### Ramp incremental tests

To assess aerobic fitness, a ramp incremental test was conducted on an electromagnetically braked cycle ergometer (Velotron; Dynafit Pro, Racer Mate, Seattle, WA, USA). After a 4-min warm-up at 50 W, a ramp of 30 W/min (for trained) or 20 W/min (for untrained) began, and the participants pedaled until volitional exhaustion. Ventilatory and gas exchange variables were measured with a metabolic cart connected to a mixing chamber (Quark CPET, Cosmed, Rome, Italy) with 10-s data averaging. The flowmeter and gas analyzers were calibrated prior to each test following manufacturer’s instructions. Heart rate was monitored and recorded continuously using a chest strap HR monitor (Polar Electro, Kempele, Finland). 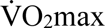 was defined as the highest 30-sec rolling average of 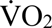.

#### Hb_mass_ and vascular volumes

Hb_mass_ was measured before and after the 4-week protocol using a modified CO rebreathing procedure, as described by (Schmidt & Prommer, 2005). Briefly, 1.0 to 1.5 mL/kg body mass (BM) of CO (determined based on sex and habitual physical activity) was injected into a closed respirometer system (Blood tec GmbH, Bayreuth, Germany) connected to a 3-L anesthetic bag filled with 100 % O_2_, and this gas mixture was rebreathed for 2 min. Before the rebreathing and 7 minutes after the start of -rebreathing, blood samples were drawn from a forearm vein. COHb and [Hb] were measured using a blood gas analyzer (ABL80 FLEX; Radiometer, Brea, CA). CO from exhaled air was measured using a hand-held CO detector (Dräeger Pac 7000, Dräeger AG, Lübeck, Germany) connected to a mouthpiece. Blood volume (BV), plasma volume (PV), and red blood cell volume (RBCV) were calculated as described by Keiser et al. (Keiser et al., 2017), using 0.91 as the correction factor for venous Hct.

### ^2^H_2_O protocol

#### Dosing and blood sampling overview

Deuterated water was administered based on lean body mass at a dose of 0.625 mL/kg LBM daily for 28 days (Oikawa et al., 2020). On Day 0, baseline blood samples and saliva were collected. Thereafter, a loading dose of 8 servings of ^2^H_2_O was provided for the participants to ingest at 90 min intervals during the day. On Days 1, 3, 7, 14, 21, and 28, the participants returned to the laboratory for fasted morning blood samples. On either day 1 or 3, additional blood samples were obtained with appropriate tubes to measure markers of iron status and a complete blood count using a clinical laboratory. The study protocol is illustrated in Figure 1.

#### Sample processing

Blood was drawn into a 2.7-mL Na^+^-citrate plastic blood collection tube (BD Vacutainer, BD, Franklin Lakes, NJ, USA) using standard procedures and centrifuged at 1500 *g* for 10 min at +4 °C. Plasma was aliquoted into tubes and frozen at −30 °C until sample preparation.

#### Hemoglobin isolation

After the initial centrifugation, 400 µl from the RBC fraction was pipetted into a 1.5 mL tube and gently washed with an equal volume of cold PBS. The solution was centrifuged at 10 000 *g* for 5 min at 4°C, and the supernatant was discarded. The remaining cells were stirred with the pipette tip and aspirated slowly several times until the suspension appeared homogenous. After repeating this procedure three times, the washed RBCs were lysed with two cycles of freezing (15 min, −80 °C, followed by thawing on ice), and then homogenized by sonication at +4 °C (20 min, 17 W). The lysed cells were centrifuged at 10 000 *g* for 15 min at +4 °C, and the Hb-containing supernatant was collected and used for HCl hydrolysis.

#### Albumin isolation

Plasma proteins were precipitated from 500 μL plasma by adding 1 mL 10% trichloroacetic acid (TCA) (wt:vol) and then centrifuging at 4500 *g* for 5 min. The supernatant was discarded, and the resultant pellet was resuspended in 500 μL ddH_2_O. Albumin was solubilized by adding 2.5 mL 1% TCA (wt:vol) in 90% ethanol and centrifuged at 4500 *g* for 5 min. The supernatant fluid was collected, and the albumin precipitated by adding 1 mL 26.8% ammonium sulfate (wt:vol). After centrifugation at 4500 *g* for 5 min, the supernatant was removed and discarded, and the resultant pellet washed once more with 1 mL 26.8% ammonium sulfate. To remove any residual free amino acids, the albumin pellet was washed twice with 1 mL 0.2 M perchloric acid. The pellet was used directly for HCl hydrolysis. The purity of both Hb and albumin fractions was confirmed by SDS-PAGE followed by transfer to a nitrocellulose membrane and Ponceau S staining (Figure 2).

**Figure 2.**
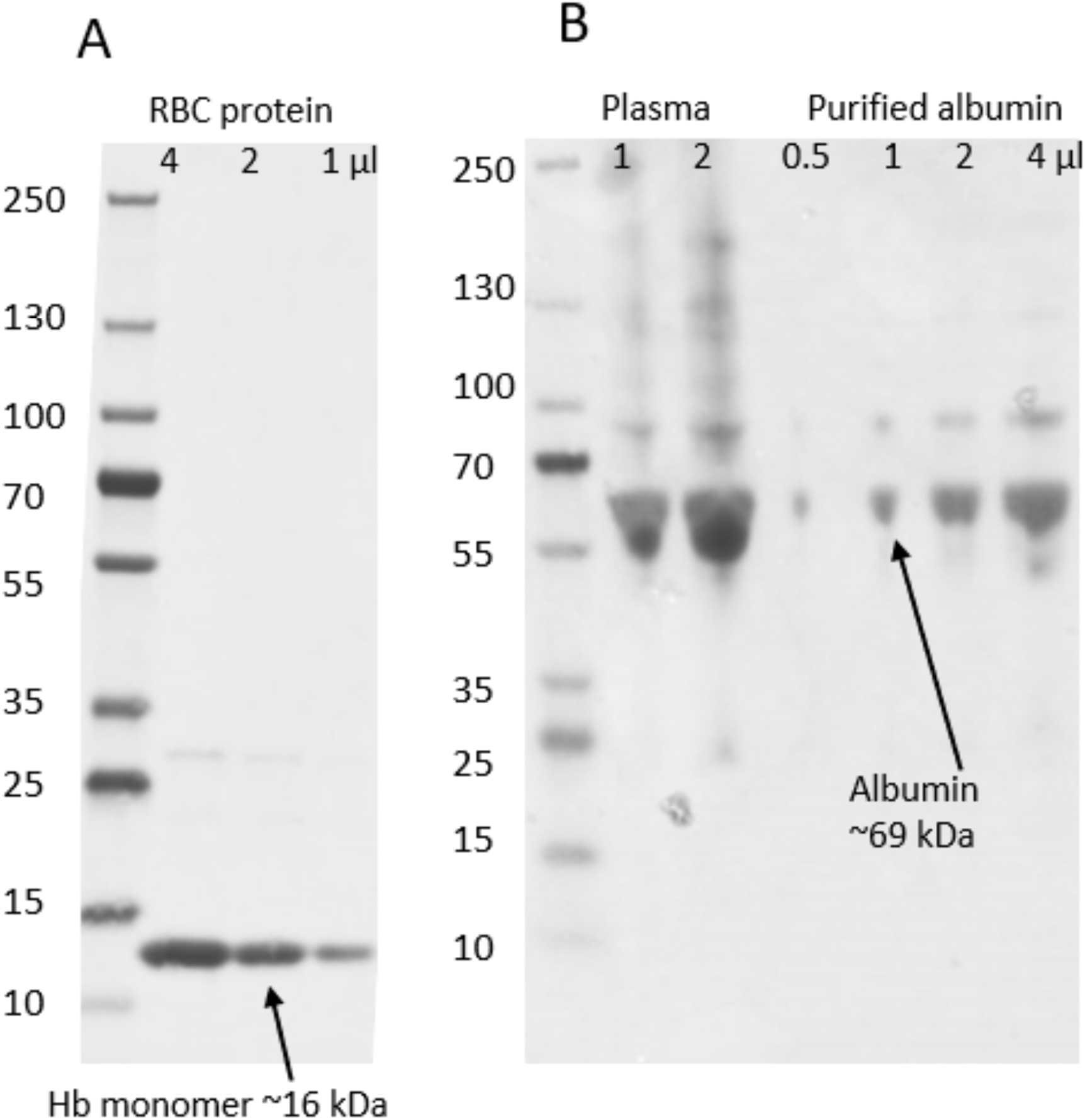
A representative image of gel electrophoresis of (A) washed and hydrolysed red blood cells compared vs. plasma, and (B) plasma vs. purified albumin from one participant. The proteins from a 4-15% TGX stain-free gel were transferred to a nitrocellulose membrane and stained with Ponceau S for imaging.

#### HCl hydrolysis and sample purification

The protein-bound amino acids in both albumin and Hb samples were liberated by HCl hydrolysis. Briefly, 2 mL of Dowex in 1 M HCl (constantly under magnetic stirring) was added to the samples, which were then vortexed and heated in an oven at 110 °C for 72 h and vortexed every 24h. Ion exchange was performed by binding the samples to columns of Dowex, preconditioned with 2M NH_4_OH and 1M HCl, then washed with H_2_O until neutral, and eluted with 2M NH_4_OH into glass tubes. The eluted samples were dried in a SpeedVac SPD1030 (Thermo Scientific, USA) at 60 °C for 18-24 h, and the resultant amino acid residue was reconstituted in 0.5M HCl. The final samples were derivatized to their N-methoxycarbonyl methyl esters and analyzed using gas chromatography combustion isotope ratio mass spectrometry by Metabolic Solutions Inc. (Nashua, NH, USA).

#### Measurement of ^2^H in body water

Saliva samples were taken independently by the participants using Salivette cotton swab tubes (Sarstedt, Nümbrecht, Germany) and frozen immediately. The frozen samples were thawed at room temperature and centrifuged at 1500 *g* for 10 minutes at 4°C to collect the fluid, which was frozen until analysis. The samples were prepared for ^2^H enrichment analysis by diluting them in 1:35 in distilled water and vortexing. ^2^H enrichment of saliva was determined by cavity ring-down spectroscopy using a liquid isotope analyser (Picarro L2130-I analyser, Picarro, Santa Clara, CA) with an automated injection system. Each saliva sample was injected six times with the mean average of the final three values taken as the final value. The coefficient of variation from the last three samples was <3%.

### Data analysis

#### Calculation of FSR and ASR

FSR is defined the rate at which ^2^H-labeled alanine is incorporated into the protein when normalized to the total abundance of the precursor pool per unit of time. FSR and ASR were calculated using the following equations:

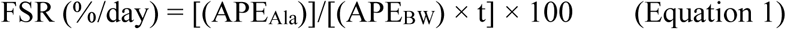

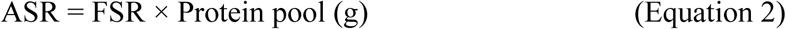

where APE_Ala_ = deuterium enrichment of protein-bound alanine, APE_BW_ = mean precursor enrichment over time, and t is the time between blood samples. A ^2^H exchange ratio of 3.7 between body water and free alanine was assumed (Wilkinson et al., 2014).

Atom percent deuterium (APD) of Hb at timepoints of blood sampling was plotted to identify the linear phase of appearance of deuterated Hb (Figure 2, A and B). The linear phase was considered to include the last 14 days of the study. The average ^2^H enrichment in body water of the first 14 days was used for the calculation of Hb FSR to account for delays between synthesis and appearance. Albumin FSR was determined from days 1 and 3 only due to the faster turnover of this protein, using the average ^2^H enrichment in body water of the same days. To obtain the number of days needed for a complete turnover of the body’s Hb stores, red blood cell lifespan was calculated as

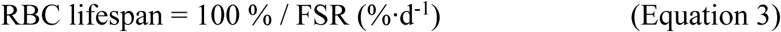

#### Statistical analysis

Means were compared using a 2-way ANOVA with training status and sex as factors. Significance was accepted at α=0.05. Normality was assessed using the Shapiro-Wilk test. Statistical analyses were conducted in IBM SPSS Statistics (version 26). Figures were created using GraphPad Prism version 10.2.

## RESULTS

Total volume of endurance training was 8.8 [3.0] h/week in trained and 1.5 [1.2] h/week in untrained during the study period (not different from self-reported baseline, P=0.78 and P=0.40, respectively) (Table 1). Relative hematological values separated by training status and sex are shown in Table 1. The trained group had a greater relative Hb_mass_, relative 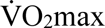 (normalized to BM and FFM), and training volume than the untrained group, but there were no differences in age, BM, FFM, absolute Hb_mass_, Hb_mass_ normalized to FFM, and [Hb] (Table 1). On a group level, no change was observed in Hb_mass_ (775 [180] g vs. 781 [202] g) or BV (5.58 [10.4] L vs. 5.53 [12.1] L) over 28 days.

Saliva APD over the 28 day study is presented in Figure 3A. Change in APD (ΔADP) of Hb-derived alanine reached it maximum between days 21 and 28; however, the ΔADP between days 14 and 21 was 97% of this maximum, which was statistically the same as the maximum, indicating a plateau (ΔADP for days 14-21: 0.063 [0.035] % and ΔADP for days 21-28: 0.065 [0.021] %, P = 0.80) (Figure 3B and 3C). Hence, the delay from Hb synthesis to appearance in blood was determined to be ∼14 days, as this timepoint marked the beginning of a linear phase in APD. Therefore, for Hb, saliva ^2^H enrichment from 14 days prior was assumed as an approximation of precursor availability at the time of synthesis.

**Figure 3.**
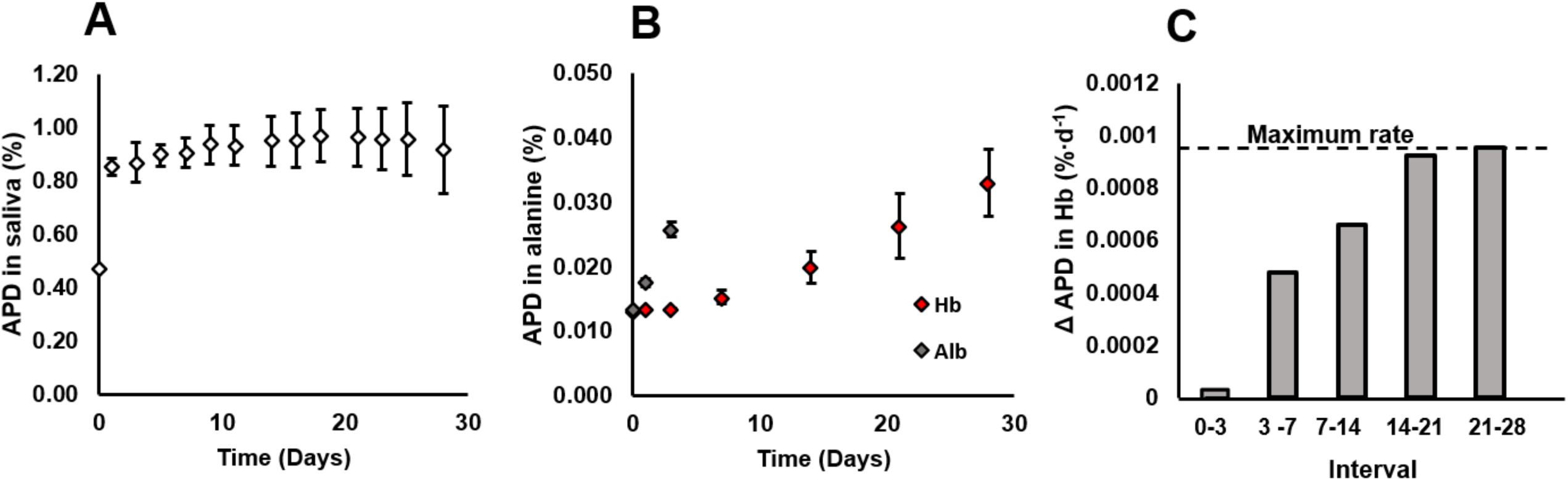
Determination of the linear phase of atom percent deuterium (APD) increase in Hb. A) APD in body water (+ SD) over the study period. B) APD in alanine (%) by sampling days in albumin (Alb) and hemoglobin (Hb). C) The rate of change in APD between sampling days for Hb. The maximum rate, measured between days 21 and 28 is shown as a dotted horizontal line. n=20 for all panels.

Values for albumin and Hb FSR are reported in Figure 4. Calculation of Hb FSR for each participant for the linear phase (days 14 to 28) resulted in a mean Hb FSR of 0.84 [21] %/day, with a range of values between 0.54%/day and 1.27%/day. These values translate to a mean lifespan of 126 [30] days, with a range of 79-187 days (Figure 3).

**Figure 4.**
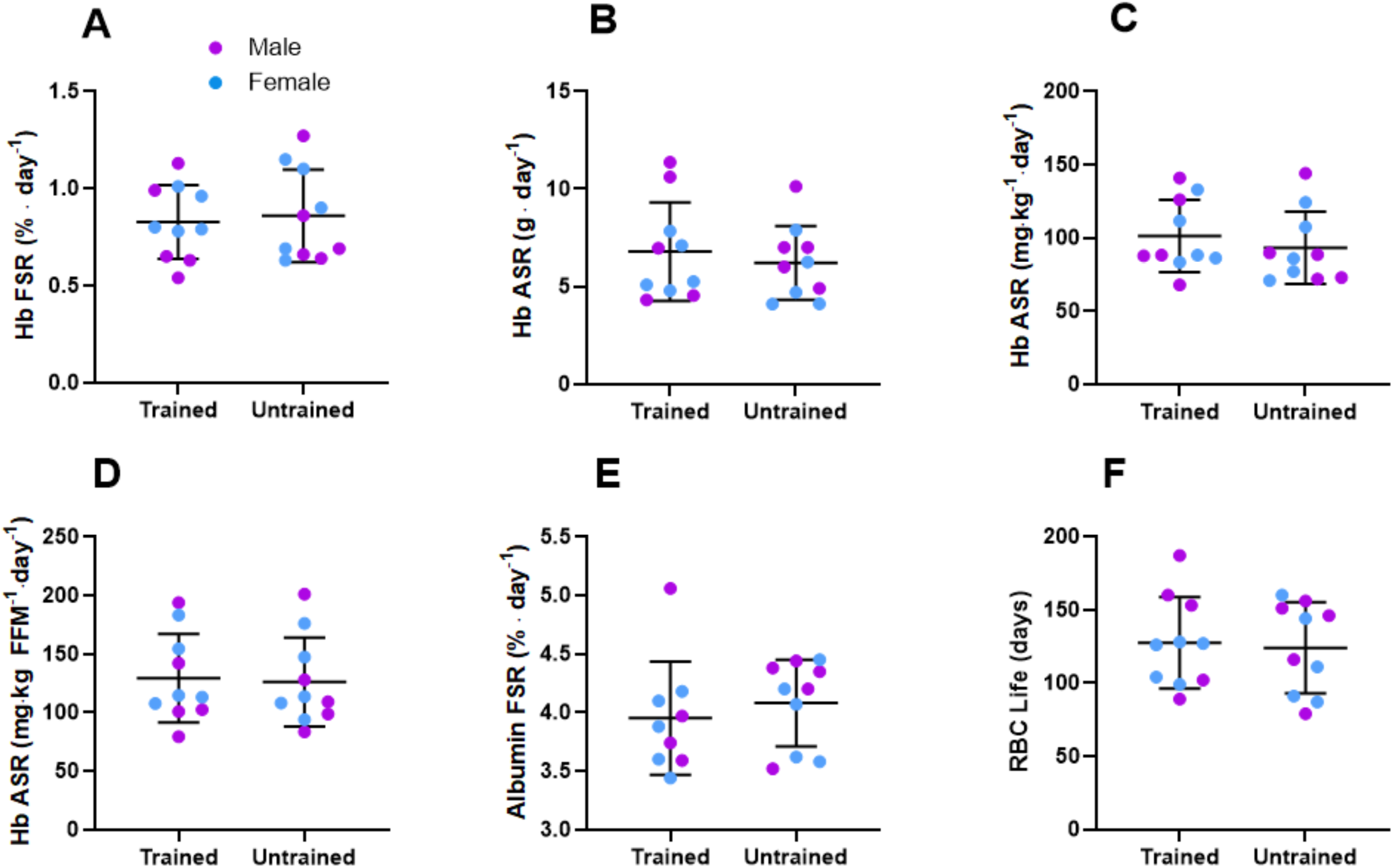
Blood protein synthetic rates for trained (n=10) and untrained (n=10) individuals. Males are represented by purple markers and females by blue markers. A) Hb fractional synthetic rate (FSR); B) Hb absolute synthetic rate (ASR); C) Hb FSR relative to body mass; D) Hb FSR relative to fat-free mass; E) Albumin FSR; F) Red blood cell lifespan calculated as an inverse of Hb FSR. The middle line represents the mean, and error bars represent one standard deviation. Individual data are shown as circles. n=20 for all panels.

As shown in Figure 4, no differences were observed between trained and untrained participants for Hb FSR (trained: 0.83 [0.19]%/day; untrained: 0.86 [0.24]%/day; P=0.74) and ASR (trained: 6.8 [2.5] g/day; untrained: 6.2 [1.9] g/day; P=0.57) values, not even when ASR was normalized to body size and expressed as mg/kg BM/day (trained: 101 [25]; untrained: 93 [25]; P=0.48) or as mg/kg FFM/day (trained: 129 [38]; untrained: 126 [38]; P = 0.85). Albumin FSR was also not different in trained (4.0 [1.3]%/day) and untrained (4.1 [0.4]%/day) participants (P=0.51).

Similarly, males and females did not differ with respect to Hb FSR (males: 0.80 [0.25]%/day; females: 0.88 [0.17]%/day; P=0.43) and albumin FSR (males: 4.1 [1.4] %/day; females: 3.9 [0.3] %/day; P=0.25). Hb ASR was also not different between sexes when examining absolute (males: 7.6 [2.6]; females: 5.7 [1.5] g/day; P=0.11), BM-normalized (males: 98 [29]; females: 97 [21] g/kg BM/day; P=0.93), or FFM-normalized values (males: 124 [43]; females: 131 [31] g/kg FFMM/day; P=0.67).

## DISCUSSION

The aim of the present study was to apply the ^2^H_2_O method for measuring protein synthesis rates to investigate the influence of training status on Hb turnover. We tested the hypothesis that habitually endurance trained individuals have a higher Hb FSR compared with their untrained counterparts due to a shorter RBC lifespan reported in this population, and a higher Hb ASR due to higher Hb_mass_ and shorter RBC lifespan. Using the ^2^H_2_O method, the mean Hb synthesis rate was 0.84%/day translating to a Hb lifespan of ∼126 days; however, in contrast to our hypothesis, we observed no effect of training status on Hb FSR or Hb ASR. Overall, the ^2^H_2_O method appears to be useful for quantifying Hb FSR and could facilitate future studies in human physiology related to erythropoiesis.

If there is a true difference in RBC lifespan between trained and untrained individuals, the lack of difference in Hb FSR between groups observed in this study is difficult to explain. A previous study reported a mean RBC lifespan of ∼70 days in trained runners (Weight et al., 1991), which is considerably lower than our measure of 127 days. Additionally, while we reported a similar lifespan in trained and untrained participants (124 days), Weight et al. reported a significantly longer lifespan for untrained participants (∼114 days) that was similar to our value. It could be speculated that our trained group was not exercising as strenuously. While we recruited endurance-trained individuals, they would not be classified as highly trained or elite athletes (McKay et al., 2022). The two groups demonstrated divergent training habits and different fitness status, as suggested by both the training load data collected during the study period as well as 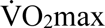 and relative hematological values (Hb_mass_, BV), which were significantly different between the two groups (Table 1); however, when normalizing hematological variables to FFM, the differences between the groups were no longer significant, perhaps indicating that the size of the relative protein pool was not substantially different in our sample. Compared with an untrained population, body mass-normalized blood volume was ∼10%, 20%, and 30% higher in moderately trained individuals, trained athletes, and elite athletes (Schmidt & Prommer, 2008), whereas in the present study, the difference was 16% in absolute units, or 10% when expressed as ml·kg FFM^-1^. This discrepancy could explain the similar Hb ASR, which would be expected to differ even with a similar Hb FSR. Whether the different methodological approaches explain the lack of differences between groups in our study is unclear.

The derived average lifespan of 126 days is similar to the canonical ∼120-day lifespan of RBC often reported (Berlin et al., 1959; de Back et al., 2014; Shemin & Rittenberg, 1946; Zhang et al., 2018) The first stable isotope experiments by Shemin and Rittenberg in the 1940s (Shemin & Rittenberg, 1945, 1946), and more recently Khera and colleagues (Khera et al., 2015) employed ingested ^15^N-glycine to determine RBC lifespan. The average RBC lifespan based on these experiments was 127 (Shemin and Rittenberg) and 106 ± 21 (Khera et al.) days. Unlike the present method, which measures synthesis rates, these methods measure the decay in isotope abundance to estimate the average lifespan. Other investigations using radioactive di-iso-propyl-fluorophosphonate (DP^32^F), ^51^Cr, or biotin labelling, or endogenous carbon monoxide production, have resulted in mean life spans of 110 to 126 days (Bentley et al., 1974; Bratteby & Wadman, 1968; Mock et al., 2011; Zhang et al., 2018). With previous studies, interindividual variation is considerable (i.e., SD > 20 days), which is congruent with the present data. Werre et al. (2004) have argued that methods utilizing cell-membrane labelling (by biotin or ^51^Cr) underestimate the true lifespan of RBC due to membrane shedding, which causes the cells to shrink in size, losing 20% of their membranes, including associated proteins, and Hb content during their lifespan. Thus, they suggest the true lifespan to be ∼150 instead of ∼120 days; however, the reduction in RBC size happens first through a reduction in water content (up to 25 days of age) and only later through loss of membrane and Hb, which becomes apparent in older cells (>80 days). Hence loss of Hb through vesiculation is unlikely to affect the present FSR measurement, as data were based on the first 28 days of Hb synthesis. Given that Hb_mass_ was stable in our participants, the synthesis and breakdown rates for Hb should be approximately equal.

To calculate protein synthesis from metabolic tracer data, a few assumptions must be made. First, we need to measure APD in the correct precursor pool. Since the measurement period in the present study was weeks rather than hours, it is almost certain that the ^2^H-labelled free alanine pool present in bone marrow was in equilibrium with ^2^H-labelled body water and therefore this assumption can be considered met. Second, we need to measure APD in the correct protein of interest. As 97% of RBCs is Hb, we analyzed proteins liberated from the RBC fraction of blood samples without further Hb-isolating methods. Third, we must assume that no recycling of the precursor takes place. While ineffective erythropoiesis (Sannolo et al., 1992), the destruction of developing RBC in the marrow, can be significant in disorders of hemoglobin synthesis (e.g. folate or iron deficiency or thalassemia), we recruited participants with no known conditions affecting erythropoiesis, meaning this process is unlikely to be a major factor contributing to precursor recycling. Significant (early) shedding of Hb-containing vesicles from circulating RBCs (Werre et al., 2004) is possible and cannot be ruled out. Finally, for Hb synthesis, because there is a delay in the appearance of labelled Hb relative to synthesis (Figure 2), we need to properly align the precursor and protein APD values for calculation. Given that it took ∼2 weeks for ^2^H label appearance in Hb to stabilize, we chose to use the body water APD from 2 weeks prior when calculating Hb FSR. As body water enrichment was relatively stable, this decision had a minor effect on our calculations.

Since plasma volume is equally increased in trained individuals, and intense exercise has been found to acutely stimulate albumin synthesis (Nagashima et al., 2000; Yang et al., 1998), it is plausible that albumin FSR would also be higher in athletes compared to non-athletes; however, based on present data, chronic endurance training does not appear to increase the turnover of albumin. We found a similar albumin FSR in our sample of trained and untrained individuals (3.95 vs. 4.08 %·day^-1^) determined from a 48-h change in albumin APD, between days 1 and 3. Our value aligned well with values previously reported in the literature (Fu & Nair, 1998; Gersovitz et al., 1980; Previs et al., 2004). The absence of elevated albumin FSR in athletes despite evidence of an acute increase in albumin FSR in response to intermittent exercise (Yang et al., 1998) might be due to the habitually trained status of our participants; these volunteers are chronically hypervolemic and thus may not respond detectably to acute training stimuli by increasing their albumin turnover; however, albumin ASR (not measured) may be higher in athletes, as would be expected due to the higher plasma volumes in this population: the trained group had on average 20% more plasma. Indeed, previous research has found an increase in albumin FSR and ASR in response to both moderate and intense exercise (Nagashima et al., 2000; Sheffield-Moore et al., 2004).

In conclusion, the novel method to quantify Hb synthetic rate by deuterated water labelling is feasible and provides results aligning with expected rates of erythropoiesis, previously only indirectly derived from RBC lifespan measurements. Contrary to our hypothesis, the Hb FSR did not differ between endurance-trained and untrained, healthy adults. Future studies may use the technique to investigate the erythropoietic process in various clinical conditions or in response to nutritional or environmental stressors, and for cross-validation of existing techniques in hematology.

## DATA AVAILABILITY

The data supporting the findings of this study are included within the figures or are available from the corresponding author upon reasonable request.

## AUTHOR CONTRIBUTIONS

All authors contributed to the conception and design of the work and the acquisition, analysis, and/or interpretation of the data. HK and MJM wrote the first draft of the manuscript, and all other authors critically revised the manuscript for intellectual content. All authors approved the final version of the manuscript and agreed to be accountable for all aspects of the work in ensuring that questions related to the accuracy and integrity of any part of the work are appropriately investigated and resolved. All persons listed as authors qualify for authorship, and all those who qualify for authorship are listed.

